# Translation rate prediction and regulatory motif discovery with multi-task learning

**DOI:** 10.1101/2022.05.03.490410

**Authors:** Weizhong Zheng, John H.C. Fong, Yuk Kei Wan, Athena H.Y. Chu, Yuanhua Huang, Alan S.L. Wong, Joshua W.K. Ho

## Abstract

Many studies have found that sequence in the 5’ untranslated regions (UTRs) impacts the translation rate of an mRNA, but the regulatory grammar that underpins this translation regulation remains elusive. Deep learning methods deployed to analyse massive sequencing datasets offer new solutions to motif discovery. However, existing works focused on extracting sequence motifs in individual datasets, which may not be generalisable to other datasets from the same cell type. We hypothesise that motifs that are genuinely involved in controlling translation rate are the ones that can be extracted from diverse datasets generated by different experimental techniques. In order to reveal more generalised cis-regulatory motifs for RNA translation, we develop a multi-task translation rate predictor, *MTtrans*, to integrate information from multiple datasets. Compared to single-task models, *MTtrans* reaches a higher prediction accuracy in all the benchmarked datasets generated by various experimental techniques. We show that features learnt in human samples are directly transferable to another dataset in yeast systems, demonstrating its robustness in identifying evolutionarily conserved sequence motifs. Furthermore, our newly generated experimental data corroborated the effect of most of the identified motifs based on *MTtrans* trained using multiple public datasets, further demonstrating the utility of *MTtrans* for discovering generalisable motifs. *MTtrans* effectively integrates biological insights from diverse experiments and allows robust extraction of translation-associated sequence motifs in 5’UTR.

## Introduction

For eukaryotic translation systems, the primary point of regulation of translation occurs at the initiation stage [17, 38, 31]. The 5’ untranslated regions (5’UTR) encode many important sequence features, collectively shaping the initiation efficiency [12, 3]. While the molecular mechanism of translation regulation has been studied for decades, it remains challenging to predict how well the 5’UTR of a transcript impacts its translation. The discovery of predictive sequence motifs in the 5’UTR and how the interplay of these elements exert their regulatory effects remain an important research area in molecular biology.

There is a rapid evolution in methods, both experimental and computational, to identify the regulatory code of translation control from the high throughput sequencing-based translation rate profiling. The most common sequencing-based techniques include Ribosome Profiling [16], Fluorescence-Activated Cell Sorting (FACS) screening coupled sequencing, and Massively Parallel Reporter Assays (MPRA) [10, 32]. Previous hypothesis-driven methods required one to firstly have a set of query elements and test them in the RP dataset or FACS-screening dataset afterwards [12, 24, 27, 8]. More recently, with the rise of deep learning methods in the field of regulatory genomics, data-driven approaches have emerged as another direction to discover the regulatory code by the model without hypothesis. [10, 1, 40, 20, 4]. MPRA methods can yield by far the highest to half millions of 5’UTR-translation rate pairs and offer rich data for the deep learning methods to capture the detailed pattern of how translation rate is controlled by the sequence context in 5’UTRs [32, 18].

However, deep neural networks can capture unintended rule that relies on dataset-specific covariates, also called the ‘short-cut’ [13, 11]. Even though seemingly successful in one dataset, short-cut features may fail to generalise to slightly different circumstances. Existing studies often only extracted and evaluated the elements in a single library or with datasets generated by the same technique. Karollus et al. have found that the sequence motifs discovered from the MPRA dataset can hardly generalize to endogenous datasets [18], which is likely caused by the difference of the truncated UTRs used in MPRA datasets from their full-long origin. This could also be explained by the fact that each of these translation rate measuring methods is probing the translation control system from a different angle. This poses a challenge to distinguish actual translation-related motifs from the false positives caused by dataset-specific artefacts [13].

To resolve the issue, we propose a multi-task translation rate prediction model *MTtrans* which can gather insight from various datasets for more accurate prediction and co-optimize a set of translation regulatory features that generalise across techniques. The prediction of TR for each dataset is taken to formulate individual tasks for *MTtrans*, which turns up to predicting highly related tasks simultaneously. Multi-task learning with the shared encoder can be regarded as a regularisation technique in our case to prevent the model from over-committing to a single task, therefore enforcing the generalisability of the learnt features [22, 34]. We here assume that the genuine regulatory elements can be found in multiple datasets, even if they were measured by different experimental techniques. The fundamental assumption of our work is that *robust sequence features* in 5’UTR that predict translation rate across multiple datasets generated by different experimental techniques are more likely to be actual regulatory elements for mRNA translation. Our shared encoder design enables the model to craft features beneficial for all tasks while retaining task specificity in the higher layers. In this work, we demonstrated that our model could outperform existing methods in both synthetic sequences and endogenous sequences. Further, our model identifies sequence features that are generalisable to an independent dataset collected from yeast. We determined what combinations of tasks would lead to the most robust predictive models and derive sequence motifs. As further validation, we conducted an independent FACS screening experiment to validate the identified sequence motifs.

## Methods

### *MTtrans* model

Our multi-task learning model *MTtrans* applies the hard parameter sharing to encode the information from several task-specific inputs. The model consists of a shared encoder *f*_*e*_ to embed sequences into a shared space and the task-specific towers 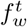 to capture the variance of the translation rate for the t-th task. Suppose a total of T tasks which are sequence libraries from any techniques, are integrated to train the model and the data for task *t* is denoted as 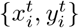 with paired translation rate 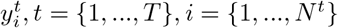.

### The shared encoder

We stacked four 1d-convolution filters layers as the main building block for the shared encoder. Raw sequences will first be one-hot encoded, ending up a 4 dimensional input 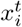 for *conv1*. Noted we don’t include any pooling operation in the encoder following the convolution layer for a better sense of the location. Instead, Batch Normalisation (BN) and drop-out operation is performed after each convolution layer to stabilise the activation and have a more robust model performance. The hidden unit at layer l is com-puted by 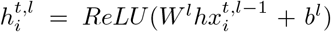 and the batch-normalized activation by 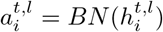.

### The task specific tower

The number of towers corresponds to the number of tasks selected. The t-th tower 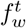 consists of 2 layer Gated Recurrent Units and an output dense layer. The hidden size of the GRU is set to 80 and the hidden state of the last time point is taken to connect to the output layer. Taken together, the translation rate is predicted by 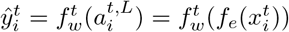 wherein 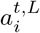 is the activation from the last convolution layer.

### Model training

One of the biggest differences in training our multi-task model is that only the t-th towers will be updated at one time. Technically, there is a task switch activating the corresponding tower for each mini-batch sampled from *χ* ^*t*^, while the shared encoder receives gradient for all the tasks. Each instance in *χ*^*t*^ is a pair of sequences 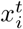 and the translation rate 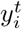 which is defined by the mean ribosome loading for MPRA tasks [32] and the translation efficiency for RP tasks [2, 15, 37]. The cost is quantified by the naive mean square error weighted by *λ*^*t*^ for *t* ∈ {1, .., *T*}:

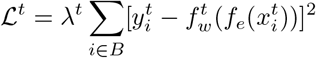

A learning rate scheduler is applied to wrap the *Adam* optimizer so 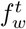 and *f*_*e*_ will be updated in a rate that is boosted in the beginning and then goes through a dramatic decay. The small learning rate in the late training phase restricts the parameters in a narrow range. Therefore, each 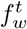 docks to a similar 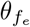 and stabilise the models. The learning rate *ϵ* is mediated by the largest dimension of the model *d*, two constants *τ*_1_, *τ*_2_ and a step-renewing variable *δ*, following by *δ* = *δ* + min(*τ*_2_, 2*δ*).

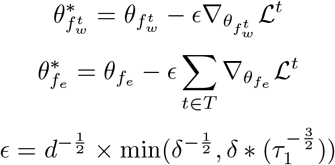

### Model evaluation

All the datasets used in our study were split into train, validation and test set. For all the MPRA tasks, the same train-test splitting was kept as in Sample et al [32] so that the model performance was directly comparable. The remaining dataset was randomly splited into training and validation set in a ratio of 9:1. The training process was terminated when the validation loss converged. For the RP tasks, the entire dataset will be splited into train, validation, test set in a ratio of 8:1:1. We set up 10 runs of experiments with different random seed each time to minimize the influence of data splitting. Similarly, the prediction accuracy was calculated by averaging the results on test set by different random seed using the Spearman correlation coefficient.

### Transfer learning

Transfer learning is a useful deep learning technique to exploit the knowledge learnt from related datasets to a new, and usually smaller, dataset [26, 28]. It could also be used to evaluate how informative are the learned features when fixing the feature extractor [39]. We relied a lot on the transferred performance to compare feature generalisability. All four convolutional layers are frozen when transferring *MTtrans* 3M to the yeast dataset. We randomly re-initialised a tower module, including 2 GRU layers and an output layer, and only updated parameters in these layers. The same MSE loss was used as the objective function. To transfer to our FACS-seq dataset for classification, in addition to the abovementioned parameter fixing and tower re-initialization, we wrapped the output neuron with a sigmoid function and changed the objective function to binary classification. For FramePooling and Optimus models, we also fixed all of their convolutional layers and re-initialised the last two dense layers.

### Data pre-processing

We collected a total of 7 public translation profiling datasets to evaluate MT-trans, including 3 massively parallel assays by polysome profiling, 1 massively parallel assay by yeast growth and 3 ribosome profiling for 3 cell lines.

The massively parallel assays MPRA-U, MPRA-H, and MPRA-V were collected from GSM3130435, GSM3130443, and GSM4084997 from the Gene Expression Omnibus (GEO), respectively. Their pre-processing are described in [32]. The Mean Ribosome Loading (MRL) is defined as the translation rate for these three libraries.

A similar technique performed on the yeast uses the growth assay to probe the translation level. Sample GSM2793752 under the accession GSE104252 is used in this study with the pre-processing described in [10] and is termed MPRA-yeast. The growth rate is taken to represent the translation rate.

For ribosome profiling data, three datasets collected from HKE293T cell line [2], PC-3 cell line [15] and muscle tissue [37] were selected because they are the benchmark datasets on which the random forest model with the handcraft features [8] was built. The ribosome profiling data were first pre-processed as in [8]. Later, to prevent data leakage, we filter the isoforms with the same sequence 100nt upstream of the start codon and keep the most abundant isoforms. The translation rate is defined as the logarithmic translation efficiency (TE), which is the division of ribosome protected fragments (RPF) reads count from the ribosome profiling to the reads count of transcripts from a paired RNA-seq sample.

For our in-house FACS-seq library, the entire UTR candidates by was used to constructed the reference. The consensus sequences were taken as the reads to map back to the reference file to quantify their count using *Bowtie2* [23]. For GSE176581 FACS-seq library, the preprocessing part is done and described as in [8].

### Task combination strategies

The performance of the multi-task model is influenced by model capacity, dataset size and the relationship among tasks [34, 19]. We thereby set the channel size of the encoder to be the same for each strategy regardless of the number of tasks grouped. The dataset size of MPRA related tasks is generally higher than the RP tasks, so each MPRA task is down-sampled to represent a comparative data size with the RP tasks. Model *3R* and *3M* are two extreme strategies tasked with datasets from the same sequencing techniques. To search for an optimal task grouping strategy with diverse tasks, we empirically tested the strategy in the following order:

1. On top of *3R*, we first introduced a sub-sampled MPRA-H dataset to have a light touch on the massively parallel assay, making a new combination *sM3R*
2. Based on the previous result, we replaced the sub-sampled MPRA-H with the full-sized MPRA-H to make a new strategy *M3R*, increasing the proportion of MPRA-tasks.
3. The strategy *2M3R* consisted of two MPRA tasks, with MPRA-V added in, and all three RP tasks. The involvement of MPRA-V here alleviates the technical imbalance on RP side.
4. Adding the MPRA-U dataset, model *3M3R* integrated all the datasets so the number of towers for the RP side and MPRA side is now equal. The direction of updating for the encoder is no more biased to the RP side.
5. We then removed the smallest task RP-muscle, to make up the new strategy *3M2R*. The task infinity of RP-muscle is also the smallest of the other five.
6. By removing the task RP-PC3, the strategy *3MR* totally consists of datasets performed in the HEK-293T cell line.

### Extraction of sequence motifs from convolutional filters

The hidden features formulated within the convolution neuron represent the local sequence combinations that are useful for predicting the translation rate. Here in this study, we use a technique called Maximum Activation Seqlet [1, 32]. This technique can reveal the short sequence segments the convolutional filters are detecting, and then these segments are used to construct the Position Weight Matrices (PWMs) [1, 32]. Here we first segment an input sequence *x*^*t*^ into several subsequences (Seqlet) 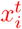 at the same length as the receptive field of the target neuron *c*. We searched for Seqlet from *X*^*t*^ that have the highest activation values for convolutional filter *c* and bagged them into *𝒫*_*c*_.

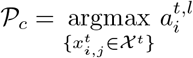

Piling up the short sequences in *𝒫*_*c*_, we can calculate the frequency of nucleotide at each position, thus is the Position Weight Matrices(PWMs). From here, the convolutional filters were converted into PWMs and were visualized by sequence logo using python package *logomaker version 0*.*8*.

### Motif similarity comparison

Important translation regulatory elements can be captured independently by models trained with different datasets. In order to identify which motifs are significantly similar to annotated RBP binding motifs, motifs similarity analysis was performed using *tomtom* from the tool-kit *meme-suite version 5*.*4*.*1*. The query motifs are explained from the models in the way we described and target motifs are labelled by RNAcompete [30] which is a built-in database in *memesuite version 5*.*4*.*1*. Only motif pairs satisfying *FDR* ≤0.05 and *E* − *value* ≤10 at the same time will be noted as significantly similar.

### Motifs matching using the extracted position weight matrices

We derived 256 PWMs from the convolutional filters. Each matrix is scanning a 9bp long sequence window, whose matching score is produced by the inner product between the flattened PWM and one-hot sequence. We scanned along the query sequence in the stride of 1 bp and we only kept the maximal matching value across windows of every position. A UTR is considered to contain the motif when this collapsed score goes above a defined threshold. Because the score distribution differs from motif to motif, we set the activation threshold to the quantiles of their score distribution.

### Building logistic regression and random forest model on PWM-derived scores

The PWMs generate a positive real number which reflects the presence and strength of the motifs when scanning the input sequence. We could obtain 256 PWM scores for each sequence, which formulate the feature set to describe a sequence. The feature vector is standard scaled across all 1,052 UTRs from the two classes of our in-house FACS-seq dataset. We implemented the model training using *scikit-learn* package version 1.1.3. The training set and test set were then split in a ratio of 9:1 with different random seeds. The performance metric F1-score and AUROC score were evaluated in the test set for the Logistic Regression (LR) and Random Forest (RF) model, as well as the three deep learning baselines.

### Identification of important motifs

Leveraging the white-box nature of the RF and LR model, we can obtain feature importance for 256 motifs and assign their regulation direction with the coefficient of LR. We first select the top half of the motifs ranking by the RF importance score. Motifs remain the same sign for all MPRA tasks that were first filtered (45 positives and 150 negatives), where we further identified 25 negative motifs by their intersection over negative LR coefficients. Because of fewer positive motifs left, we identified them from the top half of important features, resulting in 29 positive motifs.

### Design and construction of 5’ UTR library

Ribo-Seq and RNA-Seq data on HEK293T and hESC cells were extracted from RPFdb v2.0 [36] and Gene Expression Omnibus [7] respectively. Reads Per Kilobase of transcript per Million mapped reads (RPKM) was used as the primary input. Rare genes with RPKM less than 1 were eliminated. The translation efficiency (TE) score was defined as the ratio of RPKM of the ribosome protected fragments (RPFs) by ribosome profiling over the average RPKM of the transcripts by RNA-seq. Average RNA-seq RPKM was calculated from 3 HEK293T and 4 hESC samples respectively. Natural 5’ UTR sequences were defined from −100 to −1 with respect to the start codon of the coding sequence on mRNA transcripts documented in the Reference sequence (RefSeq) database [29]. The −100 to −9 sequences were extracted and attached CTAGCCACC containing a NheI overhang and Kozak sequence for cloning purposes. Synthetic sequences were generated by dividing the −100 to −9 sequences into 7mer or 31mer windows and scrambling them, followed by CTAGCCACC attachment. The natural sequences were ranked according to TE scores. Top and bottom 5% of the population in each cell line were selected and 4,800 of them were placed in the library pool, with 1,032 HEK293T specific, 1,532 hESC specific and 2,236 double specific 5’ UTRs. 3,200 synthetic sequences were also selected for the library with 1,400 generated from HEK293T specific, 1,400 generated from hESC specific and 400 generated from double specific 5’ UTRs. Together, an 8,000 library was designed. The library was then generated by the GenScript Precise Synthetic Oligo Pools service.

To insert the 5’ UTR library into an expression plasmid, the 5’ UTR sequences were firstly amplified from the Oligo Pool by Phusion DNA polymerase (New England Biolabs). The resulting fragments were digested by SbfI and BbsI (New England Biolabs) and ligated into a SbfI and NheI (New England Biolabs) digested lentiviral GFP expression plasmid using T4 DNA ligase (New England Biolabs) overnight at 16 ^°^*C*. The ligation products were transformed into chemical competent E. coli strain DH5*α* and selected by 50 *μ*g mL1 carbenicillin. The resulting library was extracted and purified using Plasmid Midi (Qiagen) kits.

### Validation of discovered motifs with the FACS screening library

Our core assumption is that collectively training the model with datasets from diverse techniques can lead to a more biologically relevant feature set. The FACS sequencing method was used here as a complement of the RP and MPRA to test the robustness of the sequences motifs derived from *MTtrans*. When sorting the cells carrying 5’UTRs with high translation efficiency, their reads count *y* in the sorted bin becomes a proxy of the translation efficiency.

We firstly encoded the 5’UTRs by *MTtrans* to extract the feature map of layer 3. Then we searched the sequences that can uniquely activate the motifs. For each convolutional filter, the 5’UTR sequences were then ranked by the activated values in the feature map. The top-ranked sequences were regarded as the activated sequence for the corresponding filter. Each sequence can only be assigned to one filter. When one sequence ranks top for multiple filters, the filters with the highest activation values will be assigned to it, and it will be added to the uniquely activating set for each convolution filter (channel). The effect of a motif converted from filter *c* is therefore defined as the mean of log count in the uniquely activating set minus the the average read count of the library. In other words, if the reads count of the activated set is higher than average, we assign it as a positive motif and *vice versa* for negative motif. Lastly, we compared the estimated effect with the effect they displayed in the FACS library. If their direction is the same, the motif effect is consistent in the new library.

### Cell culture and cell sorting

HEK293T cells were obtained from the American Type Culture Collection (ATCC) and cultured in DMEM supplemented with 10% FBS and 1× antibiotic–antimycotic (Life Technologies) at 37 °C with 5% CO_2_.

Lentivirus production of the 5’ UTR library was similar as previously described [9]. The resulting lentivirus was titrated to a multiplicity of infection of 0.15 to achieve low copy number delivery of the 5 ‘UTR library into HEK293T cells while achieving 100 fold representation.

On day 7 post-transduction, cell sorting was performed on a BD Influx cell sorter (BD Biosciences). Live single cells were gated and sorted by GFP signals. A two-round sorting approach was used by enriching the total GFP population and further sorting them into three bins according to their GFP intensity.

### Deep sequencing

Genomic DNA of collected samples were extracted by DNeasy Blood and Tissue kit (Qiagen). The inserted 5 ‘UTR region was amplified by PCR using Kapa HiFi Hotstart Ready-mix (Kapa Biosystems). The fragments were gel purified by QIAquick Gel Extraction Kit (Qiagen) and sent to HKU LKS Faculty of Medicine Centre of PanorOmic Science for SMRTbell Library construction and Single Molecule, Real-Time (SMRT) sequencing (PacBio).

## Results

### MTtrans learns the shared patterns from multiple experimental systems

*MTtrans* is developed as a multi-task learning method to account for the variance in translation rate given by different experimental systems. We adapted the canonical hard parameter sharing architecture which consists of a shared encoder (bottom) and several task-specific upper layers (towers) (Figure 1). To train the model, we collected datasets from different translation systems and diverse techniques, including three artificially synthesised libraries constructed with MPRA combined with polysome profiling in human HEK293T cell line (MPRA-U, MPRA-H, MPRA-V), one synthetic library from MPRA with yeast growth as the readout (MPRA-Yeast) and three ribosome profiling studies measuring the translation efficiency of the nature transcripts from HEK293T cell line, PC-3 cell line and human muscle tissue respectively (RP-293T, RP-PC3, RP-Muscle).

**Fig. 1.**
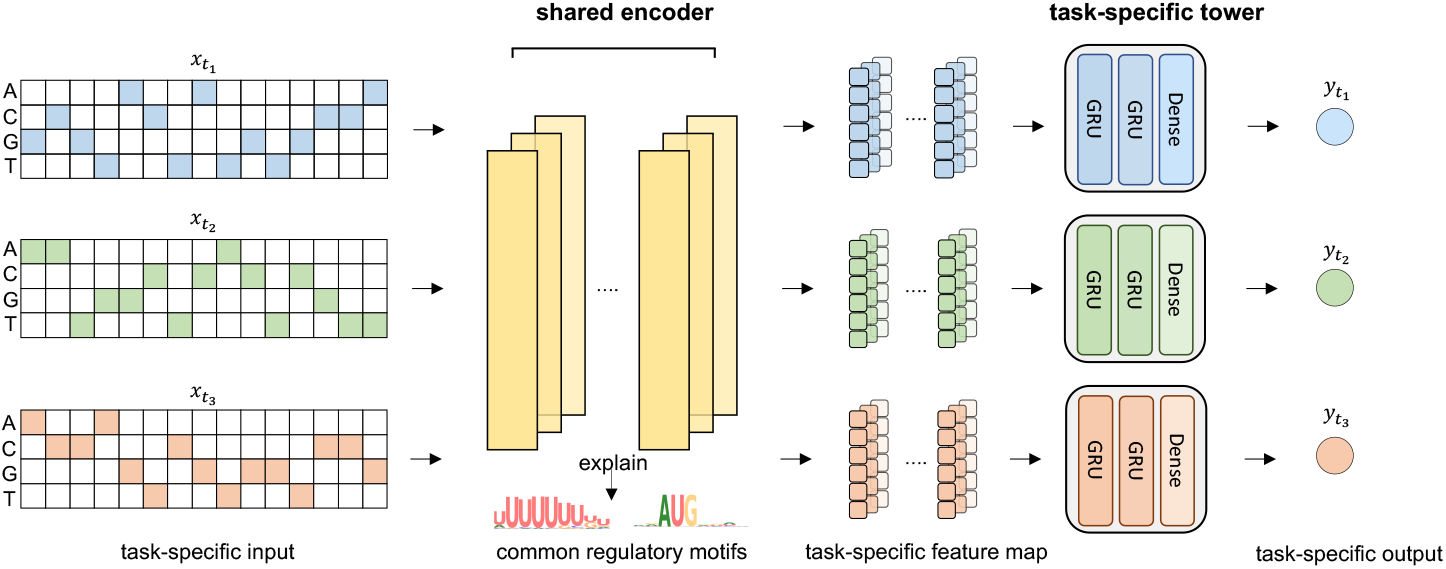
The model architecture of our multi-task learning method *MTtrans*. The model consists of a shared encoder and several task-specific towers. We use the shared CNN encoder (bright yellow) to extract features and transform task-specific input *x*_*t*_ into task-specific feature map. The task specific tower takes in the feature map and predicts the translation rate *y*_*t*_ for the corresponding task. Blue, green and orange color stand for different tasks.

In general, we break down the typical translation rate prediction process into two parts: sequence feature extraction and regression. We let the shared bottom take care of sequence feature extraction. Basically, the bottom module was mainly made up of four 1D convolution layers, analogous to the translation initiation process of scanning through the input sequences from the 5’ to 3’ end to detect the patterns of interest. One important merit of using convolutional neural networks is the emergence of many network explanation approaches in recent years [4, 14, 33], allowing for the visualisation of the regulatory signal encoded in the higher layers. The towers will learn to reorient the extracted pattern for each task and make the regression from sequence pattern to translation rate (Figure 1). The tower module is built up with a two-layer Gated Recurrent Unit network (GRU) to deal with the various length of inputs and a dense layer to project the GRU memory to the output value, which is the predicted translation rate.

### *MTtrans* better coordinates MPRA tasks and improves translation rate prediction

We started with the MPRA datasets, in which the translation rate is measured with mean ribosome loading (MRL), to evaluate the effectiveness of our *MTtrans* model. We included two published model *OptimusN* [32] and *FramePooling* [18] for comparison. Another baseline called *mixing* was trained with a dataset merging all the MPRA libraries. The involvement of *mixing* was to account for the benefit of being trained from a larger integrated dataset.

Our multi-task method can predict the translation rate more accurately than the alternatives in all MPRA tasks, regardless of input length or sequence origin (Fig.2 a&b). In the largest MPRA dataset (fixed length random UTRs, short for MPRA-U), our model can consistently exceed the state-of-art result (*r*^2^ = 0.932 by OptimusN, *r*^2^ = 0.928 by FramePooling) [32, 18] and push translation rate prediction to the highest of 0.947 in *r*^2^ (*p*=0.0037 compared to FramePooling, one-sided unpaired T-test, Figure 2c & Supplementary Table 2). Interestingly, model *mixing* showed a performance loss on the fixed length human UTRs (task *MPRA-H* for *MTtrans*, the left panel of Figure 2b), suggesting that it could be harmful to simply mix the datasets probably due to covariates like batch effect and shift in sequence composition between randomly synthetic sequences and natural sequences. When modelling 5’UTR sequences with varying lengths, it turned out that using more data together with GRU layers can boost the performance strikingly. Thus, with the designed task-specific towers, our method could alleviate the conflict and could effectively coordinate datasets with different study designs to finally benefit all the tasks.

**Fig. 2.**
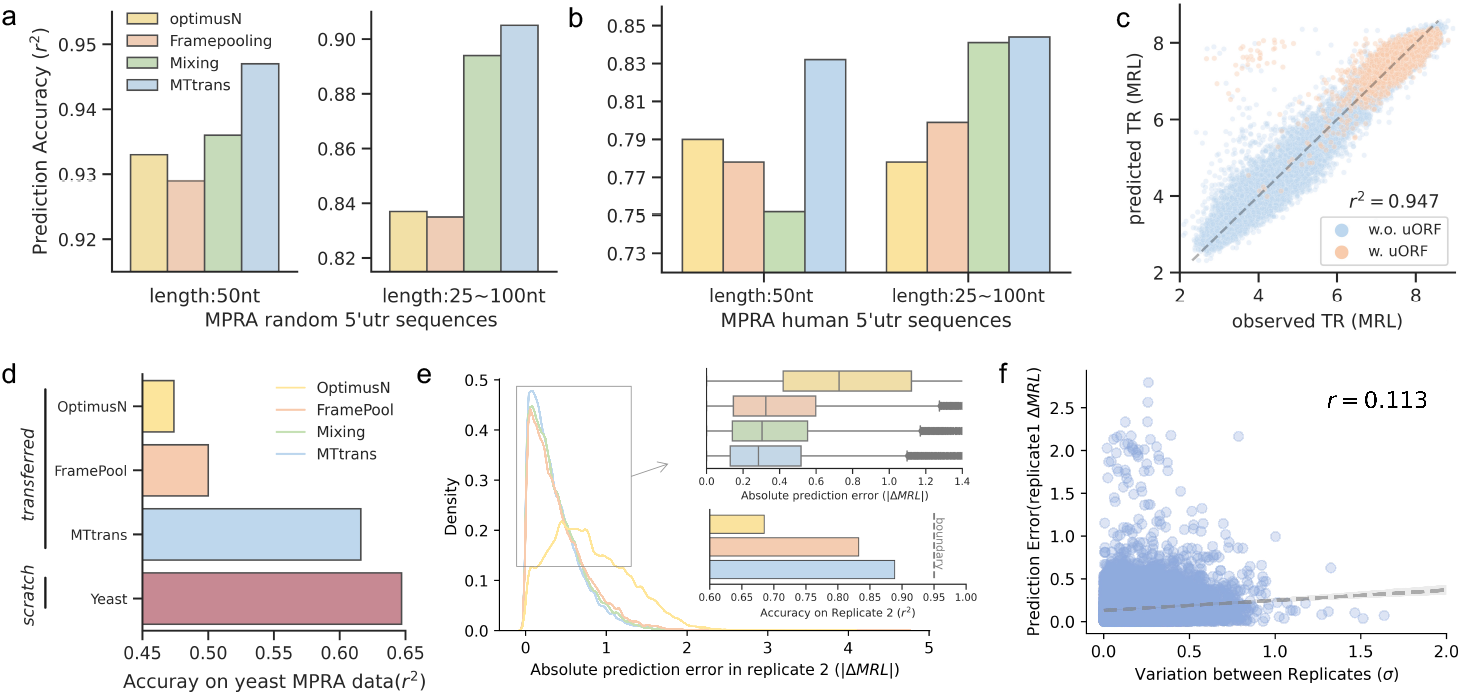
MTtrans make accurate and robust translation rate prediction by combining the signal from different task. **a** Performance for predicting mean ribosome loading (MRL) on fixed length random synthetic sequences MPRA dataset (left) and also a varying length synthetic sequences dataset(right). These 2 dataset forms 2 MTtrans prediction tasks which were learned simultaneously.. Prediction performance was measured by the *r*^2^ between the observed mrl value and the predicted in the held-out test set (n=20,000 for fixed length and n=7,600 for varing length). OptimusN [32], FramePooling [18] and the counterpart MTtrans model simply trained by mixing all the dataset were compared by evaluating them on the same test set. **b** Performance for predicting mean ribosome loading (MRL) on the MPRA dataset ofs human 5’utr sequences (left) and also a varying length synthetic sequences dataset(right). Prediction performance was measured by the *r*^2^ between the observed mrl value and the predicted in the held-out test set (n=25,000 for fixed length and n=7,600 varing length). OptimusN [32], FramePooling [18] and the counterpart MTtrans model simply trained by mixing all the dataset were compared by evaluating them on the same test set. **c** the scatter plot showing the performance for task fixed length random UTRs (MPRA-U). A high fidelity of *r*^2^ at 0.947 is reached. Sequences with uORF was colored with blue and sequences without uORF with orange respectively. **d** Transferring encoder trained from human MPRA datasets to MPRA-Yeast dataset. All the parameters in encoder are fixed for *OptimusN, FramePool* and *MTtrans*. While for model *Yeast*, the entire model is trained *from scratch* in MPRA-Yeast. **e** Robustness shown by the absolute prediction error in technical replicate. Absolute prediction error is calculated by subtracting the model prediction for MPRA-task to the label from MPRA-U replicate-2. The larger density plot showed the overall spread of error and the value ranging from 0 to 1.4 |*ΔMRL*| magnified in the upper right panel. The upper right box plot highlighting the densest region to display the mean and quantile of the error distribution. The bottom right bar plot label the consistency of model prediction for replicate-1 and observed value of replicate-2 by *r*^2^ **f** The absolute prediction error is weakly correlated to with the variation observed between two replicates. Prediction error is calculated in replicate-1. Variation between replicates is calculated by firstly normalizing MRL to *σ* = 1 for each replicate and then the absolute different of the two normalized MRL so that the dift of MRL by a unit of *σ*.

### *MTtrans* learns more transferable sequence features

To test whether the shared encoder of *MTtrans* captures a more universal feature set, we fixed the encoder and then transferred the model to a new MPRA random 5’UTR dataset (MPRA-Yeast) generated in *Saccharomyces cerevisiae*. The performance of the transferred model with the fixed encoder can imply how general these learned features [39].

*MTtrans* has learnt more transferable motifs than the single-task models (*OptimusN, FramePooling*) significantly. However, *MTtrans* was not on par with the model trained *from scratch* (box coloured with mauve in Fig. 2d), which showed the evolutionary divergence between the two translation systems despite a considerable convergence they shares. This performance gap may imply the existence of yeast-specific patterns shaped by the evolution process that can not be filled by learning more human data. The result may suggest that *MTtrans* has captured more generalisable sequence features in 5’UTR for translation initiation. Although not comparable with the yeast-oriented model, there is a significant improvement over the single-task models. Apart from seeing more sequence composition during training, using more datasets may also prevent the encoder from capturing dataset-specific patterns as it will hamper the other tasks and increase the overall loss during optimization. Overall, using the shared encoder to gather different tasks is a good strategy for learning more transferable features and may lead to genuine regulatory motifs.

### *MTtrans* is robust across replicates

*MTtrans* has achieved amazing accuracy on the MPRA-U task, but it’s reasonable to question whether it is over-fitted on the MPRA-U task and captured some unwanted batch effect. To answer that, we collected sequences having two experimental replicates from MPRA-U. Models fitted on replicate-1 were then evaluated in the test set of replicate-2.

The translation rate predicted by *MTtrans* is more consistent with that measured in replicate-2 (Fig.2 e). The absolutes prediction error was calculated between the predicted MRL and the observed MRL in replicate-2. The error made by our methods mostly fell in 1 ± *MRL* and has the closest mean of error getting to 0. The consistency, measured by the *r*^2^ in replicate-2, also showed the advantage of our multi-task method over the single-task methods. Interestingly, despite a higher accuracy than *FramePooling* in replicate-1 (Fig.2 a), *OptmiusN* made more error than *FramePooling* in replicate-2. We further asked whether the failure cases by *MTtrans* were caused by the incorrect signal in the data. We correlated the error in replicate-1 with the variance of measurement and found a weak correlation. There are sequences strongly deviated from our prediction but highly consistent between replicates, serving as interesting cases for further investigation. Overall, training with multiple datasets can correct the model to become more robust between replicates.

### *MTtrans* better predicts the translation rate of endogenous transcripts in human cell lines

We next turned to the human endogenous 5’UTRs in their natural genomic context, whose translation rate is typically assessed via ribosome profiling (RP). Due to their smaller data size and intrinsic confounders, such as regulation that occurred in the CDS region and 3’UTR region of the same transcript [24], It is naturally a harder task to predict the translation rate for RP datasets. Here-with MTtrans, we attempt to tackle this long-standing issue by adding an extra layer of features learned from MPRA datasets, whose sequence context is more revealing of translation initiation signal.

To train the *MTtrans* on ribosome profiling data, we selected three sequencing libraries from human HEK293T cell line [2], PC-3 cell line [15] and muscle tissue [37]. With the same datasets, Cao et al. built a random forest regressor (which is named 5’UTR RF in the later text) to predict the translation efficiency (TE) of each transcript [8]. They manually crafted features to describe the known regulators in 5’UTR and thousands of kmer frequencies to cover other possible elements. Their RF model and the single-task model were taken as the baseline. Highly similar transcripts of the same gene were filtered for all the datasets to avoid information leakage from the training set to the evaluation set (see Methods).

*MTtrans* greatly surpasses other methods in the noisy RP data (Figure 3a& b). Deep learning-based algorithms seem to be more suitable in modelling RP when we compare *single-task* with *5’UTR RF*, which may be explained by the learnt regulatory features not included in the RF model. A consistent performance gain was conferred by *MTtrans* in all cell lines. The improvement was not only observed in small source tasks like *muscle* but also held true for tasks of larger size (*i*.*e. PC-3*, Figure 3a). Notably, adding two human 5’UTR MPRA datasets further improved the performance of all three RP tasks, suggesting beneficial information sharing from the MPRA sequence features.

**Fig. 3.**
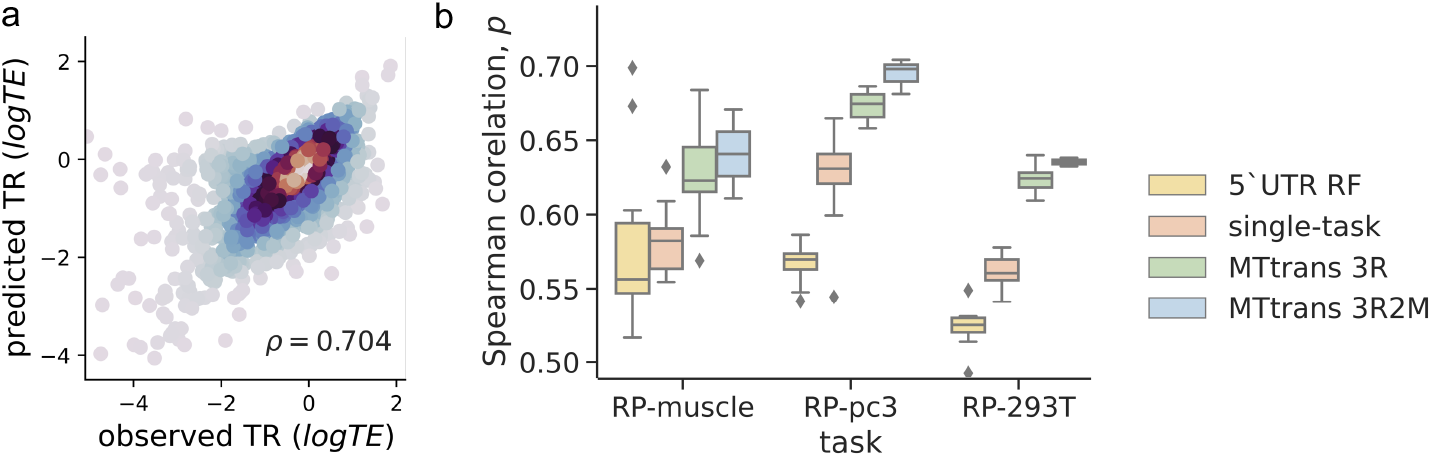
MTtrans supports a flexible task combination strategy to make effective information sharing and promote the prediction for the noisy ribosome profiling data. **a** Observed log translation Efficiency (TE) against the log TE predicted by *MTtrans* 3R on the cell line PC-3 dataset. Color shows an increasing dot density from blue to red. **b** Model performance in three ribosome profiling datasets. Model 5’UTR RF was from the The *MTtrans 3R* here was trained by the three RP datasets. The red box indicates a single-task counterpart. The blue box indicates the strategy *2M3R*. The *MTtrans* model was trained with two MPRA tasks and three RP tasks.

### Proper task selection enables *MTtrans* to balance across datasets generated from different techniques

Multi-task learning can be deleterious when there are outlier tasks sharing conflict optimising signal.[41]. To learn a set of general features, various task combination strategies were designed with a different mixture of MPRA tasks and RP tasks. To order the combination, we defined a gradient with *3R* and *3M* at the two ends, between which are a mixture at different levels (Supplementary Table 1).

Generally, task performance positively correlates with the proportion of similar tasks in the training data, but an appropriate task selection can overcome the data imbalance. For the encoder with limited model capacity, the hidden features were updated in a competitive way so that only those universal features could be kept during the training. With more MPRA tasks joined, MPRA tasks enjoyed a stable increase in the prediction accuracy, which climbed to the peak at *3M*. Interestingly, RP tasks could also benefit from the integrative training with MPRA tasks by referring to more refined sequence features identified from the large sequence space (see RP tasks in model *M3R* and *2M3R*). For model *3MR*, MPRA tasks dominate the overall tasks, and RP-293T becomes an outlier, pulling down not only RP-293T itself but also the MPRA tasks if compared to *3M*.

Thus, training the multi-task model required a careful balance of their covariates, such as the sequencing techniques and cell type. Taken together, model *3M3R* stood out from all other tested combinations (Supplementary Table 1) because it retained a comparable performance on diverse tasks regarding the technique and cell type, maximising the chance to learn more general motifs.

### Discovery of 5’UTR sequence motifs from the deeper layer of shared encoder

We next explained the convolutional filters in the shared encoder to see what features *MTtrans* has learned. There are studies that only explain the parameters of the first convolution layer [1, 40], but one could argue that features extracted from higher layers confer more abstraction and are more likely to represent the regulatory grammar [10, 4].

By taking the feature map from each layer for fitting random forest models, we found out that the deeper layers confers more information. (Supplementary Figure 2a). To explain the sequence motifs, we use the maximum activation seqlet [1, 32] to visualise individual convolutional filters at the last layer, for a longer receptive field to cover the 5 to 7bp long annotated binding sites of RNA-binding Proteins (RBP). The sequence segment that activates the neuron the most is clipped out from the sequence and grouped to generate the position weight matrix. With this method, each convolutional filter will be interpreted as a sequence motif with a receptive field of nine nucleotides. The correlation between neural activities and the translation rate can indicate the effect of the motif it derives.

The sequence features derived from the last layer of *MTtrans 3M* can reconstruct many biologically meaningful motifs. 43 motifs are significantly similar to the RBP binding motifs annotated by RNAcompete [30], indicating that *MT-trans* captured the biological meaningful elements to some degree(Figure 4). For example, HuR have been proved to bind to 5’UTR; PABR4 has high poly-U affinity [6]; RMB4 is known to suppress the cap-dependent initiation process [25].

**Fig. 4.**
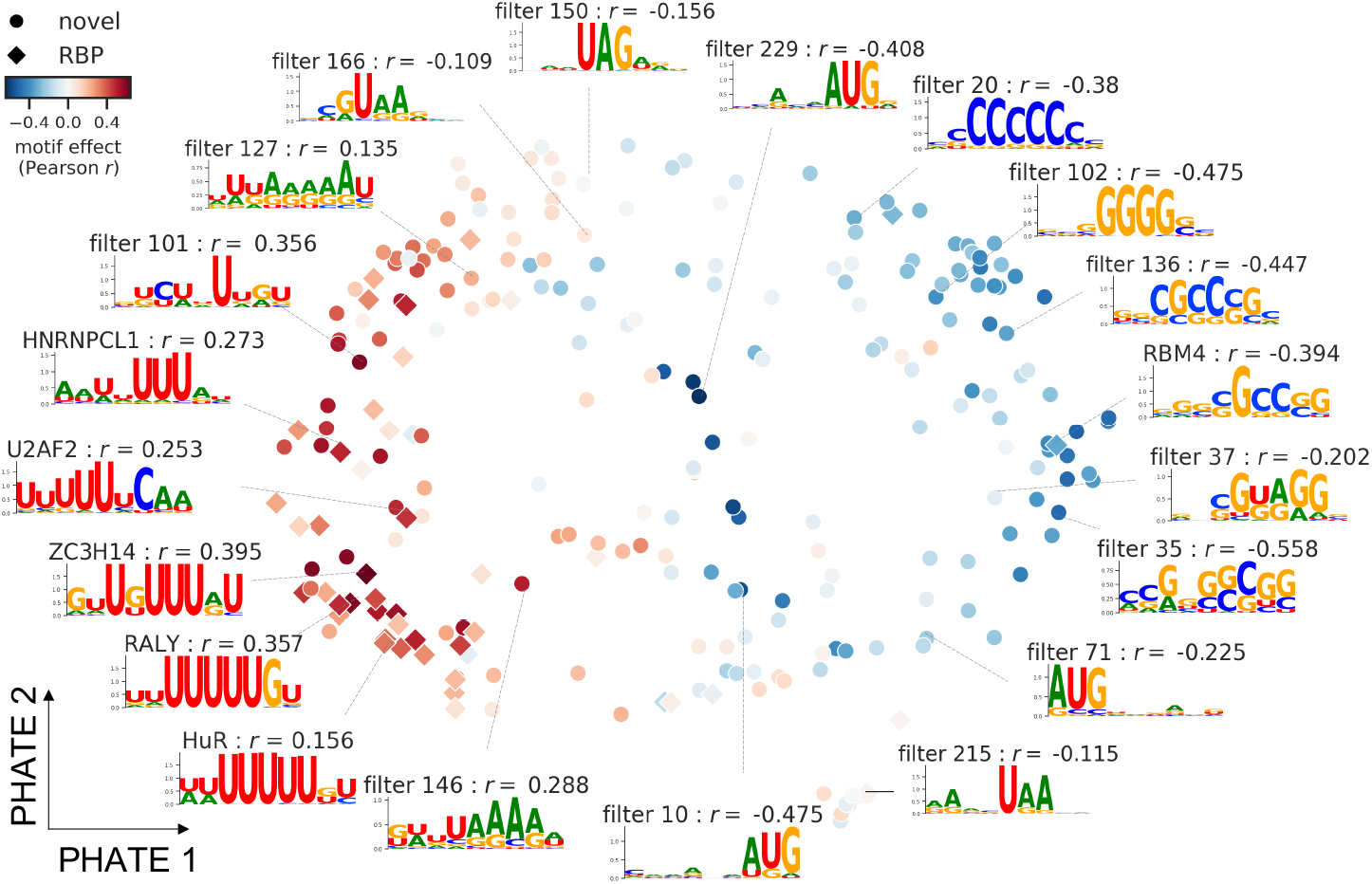
Sequence motifs discovered by MTtrans. A total of 256 motifs are projected into PHATE space by reducing the last layer feature map input from all sequences to 2 dimension. The sequence logos of convolutional filters were generated from the position weight matrices using Maximal Activation Seqlet. Colour fo the dots denotes the regulatory effect estimated in dataset MPRA-H, which is the Pearson correlation between motifs activity and translation rate. *Diamonds* indicate motifs that are significantly similar to known RBP in [30].

The upstream start codon (uAUG) and the Kozak consensus are well-characterised regulatory signals that can alter the translation initiation [17, 38, 21]. AUG is present in a substantial fraction of the found motifs. 56 of the 67 uAUG detecting motifs, the majority of which include guanine right to the uAUG, are negatively correlated to the translation rate. The positive uAUG motifs, on the other hand, represent a more interesting feature set as they define the sequence context to cancel the upstream initiation. An apparent pattern is a Uracil or Adenine flanking around the uAUG, which violates the preference of the Kozak consensus. MTtrans also re-discovered upstream stop codon UGA and UAA in filter 150 and filter 215.

In conclusion, by interpreting the learnt shared encoder, we can rediscover many genuine regulatory signals, suggesting the advantage of *MTtrans* as a motif discovery tool.

### The discovered regulatory motifs can be experimentally validated

To validate whether the discovered motifs remain predictive in datasets generated by a different experimental technology, we generated an in-house library with FACS-seq to measure the translation rate of a newly designed 5’UTRs library. 8,000 UTR sequences of length 99bp were inserted into vectors to control the translation of the GFP protein. The engineered plasmids were then delivered into the HE293T cell line, which is the same as the MPRA datasets that *MT-trans* was trained from. Finally, we applied FACS screening to sort cells by their GFP fluorescence and collected highly fluorescent cells for further sequencing so that the read count could represent the translation rate of the 5’UTR sequence (Figure 5a). When using neuron activation to detect the presence of motifs in a sequence, 72 motifs (out of 123 positive motifs) showed a higher read count than the average in the FACS library. For negative motifs, 83 out of the 141 motifs showed a lower read count in this dataset. The result suggested a (*p*=0.0066, binomial test with *k* = 153 *n* = 256, Supplementary Figure 6b).

**Fig. 5.**
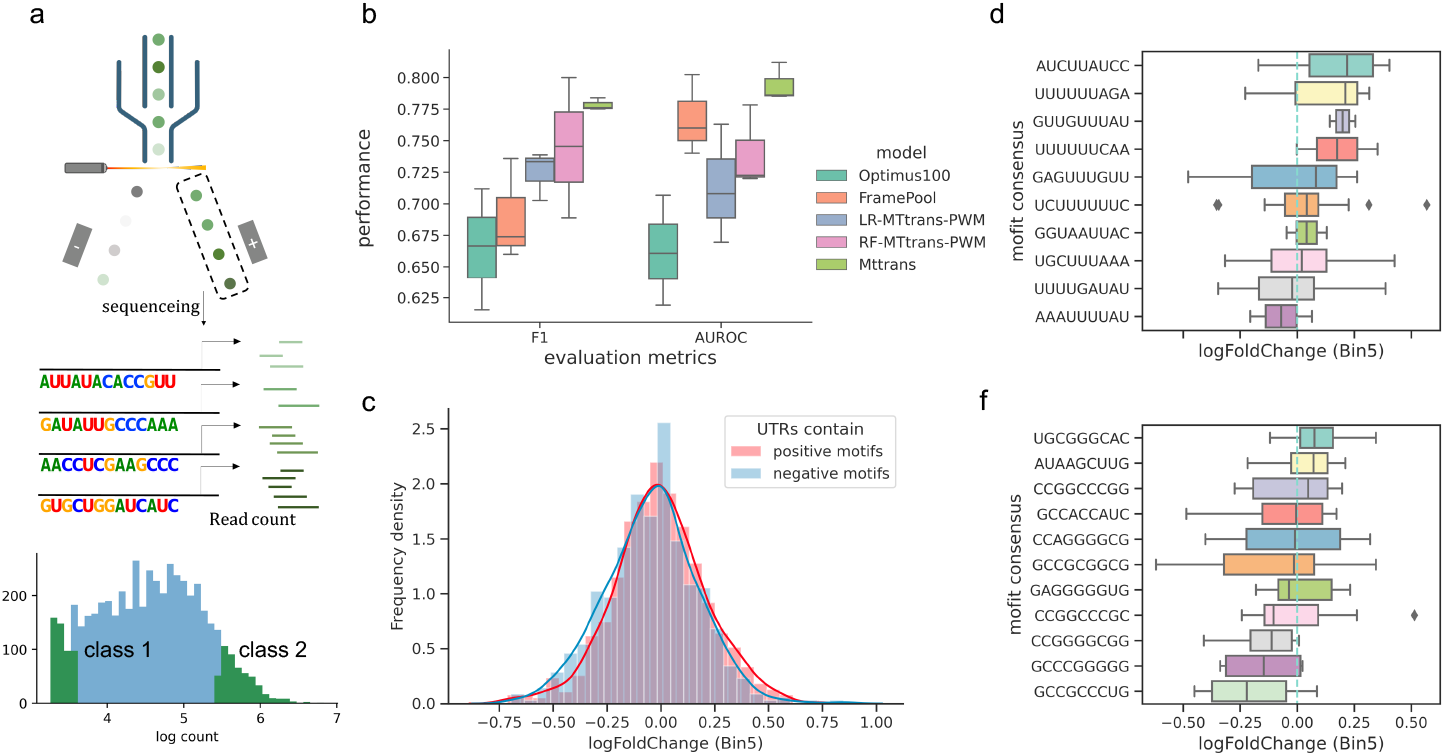
FACS experiment to validate the discovered motifs. **a** cells carrying designed UTR are sorted by their GFP fluorescence, screened for high GFP expression and then sequenced to identify the delivered UTRs. The read count of the UTRs can thus approximate its translation rate. UTRs with the lowest translation rate were grouped as the negative class (class 1) and the positive class (class 2) for the highest one. **b** Result of binary classification for the above two classes. The box shows the performance of models for different train-test splitting random seeds. LR is the logistic regression built on the features scored by MTtrans-explained PWMs. RF stands for the random forest model built using the same features. **c** The distribution of translation rate (log fold-change in Bin5) for UTRs on another FACS-seq library GSE176581. Red curve and bins, UTRs that enriched with the 29 identified positive motifs, has higher translation rate than the blue curve and bins that contains the 25 negative motif sets (*p*-value = 7.65 *×* 10^4^ for threshold of 90% quantile, one-side unpaired T-test). **d** The translation rate of sequences from GSE176581 that contain only one motif among the selected positive feature set. **f** The translation rate of the sequence from GSE176581 that contains only one motif among the selected negative feature set.

To test the extent to which the discovered motifs are preditive without the specific neural network, we used the PWMs generated by the Maximum Activation Seqlet to score the strength of motifs and perform binary classification by logistic regression and random forest for this FACS-seq dataset (Figure 5b). Using MTtrans motifs, LR and RF model succeeded in separating the two classes (average F1 0.714 & 0.740, average AUROC 0.725 & 0.745, respectively) and outperformed the deep neural network baselines *Optimus100* and *FramePool100* in F1-score. These results showed that the discovered motifs alone are robust and highly informative in modelling translation regulation.

Inspecting the learned parameters of logistic regression and random forest can give us insights into the role of each motif in the FACS-seq dataset. Basically, we can assign a motif to a positively regulating signal or negatively regulating signal according to the sign of its coefficients in the logistic regression model. Compared with the MPRA-V dataset, which also has 99bp-long UTRs, 59% of the motifs remain in a consistent regulatory direction, and it is significantly different from random (*p*-value = 0.016 by scrambling the motifs 1000 times see Supplementary Figure 8a). The feature importance scores by random forest evaluated the motifs from another way and were weighted more onto positive motifs (Supplementary Figure 8b). Combining insights from the LR and RF models, we identified 29 positive and 25 negative important and consistent motifs.

We next tested these identified motifs in another independent FACS-seq dataset to evaluate their robustness. We curated another public FACS-seq library GSE176581 [8] in which the translation rate is also represented by the enrichment of UTR-carrying cells in the high GFP fluorescence bin (log fold change of Bin5). For motif detection, a motif is considered to occur in the sequence when its PWM matching score is higher than the defined quantile threshold of the score distribution (see Method). Collectively, UTRs which score higher averagely on the positive motif set are more translated than that on the negative motifs (Figure 5c, *p*-value = 7.65 × 10^4^ for the threshold of 90% quantile, one-sided unpaired T-test), and the significance also holds for different threshold (Supplementary Figure 6), indicating a robust regulatory direction of these motif features. For individual motifs, we specifically checked out sequences that only contained one motif and with a low average score on the opponent set (Figure 5d& f). Only 10 positive and 11 negative motifs left meet the above criteria, and the majority of them maintained the same regulatory effect on the translation rate.

Overall, using two FACS-seq datasets, we demonstrated that the *MTtrans*-derived motifs are transferable to a new experimental system and can distinguish between low TR class and high TR class without the neural network. Two subsets of features, identified by PWM-based models, show a robust regulatory direction across two FACS-seq libraries. These results supported that *MTtrans* can indeed combine different datasets to yield biologically meaningful motifs.

## Discussion

In this study, we demonstrate that our multi-task learning model *MTtrans*, is effective to predict mRNA translation rate using 5’UTR sequences for diverse experimental data. *MTtrans* works by treating each dataset as an individual task so that useful information across tasks can be underlined repeatedly. Importantly, we found that this framework enables the extraction of highly robust sequence motifs that predict the increase or decrease of translation rate. This is achieved by the hard-sharing architecture in our multi-task model.

Previous studies on transcription factor motif discovery usually can extract predictive features from the first convolutional layer [1, 40]. Koo et al. even propose an ambiguous pooling to enforce the motifs detection finished in the first layer [20]. Interestingly, in the context of translation rate prediction based on 5’UTR, it is necessary to look beyond the first layer to extract features. Furthermore, we found that the improvement does not result from the longer receptive field (Supplementary Figure 2). As the deeper features are assembled from the lower layer patterns by convolution filters, translation rate prediction may thereby require better arranging the basal motifs to sense the sequence context. This might partially explain how the CNN model surpasses the *k*-mer-linear model [32] because *k*-mer models lose the sequential relationship of the features despite a broader *k*-mer coverage. Future work should explore how to best extract features from different layers of neural networks for the prediction of transcription factor binding and translation rate.

A core premise of our work is that more robust and transferable features can be extracted from a model learned from diverse data sets generated from different experimental techniques and cell types. To evaluate this hypothesis, we tested the use of different combinations of data as the tasks in *MTtrans* and found, indeed, the most cross-platform model is the one with a balanced task source (*3M3R* in Supplementary Table 1). Models trained with the data generated by the same technique, such as MPRA, obtained the specificity for the same experimental setup (Figure 2a), but fail to predict results generated by another experimental technique (Supplementary Figure 1). This suggests that such models are learning the short-cut to some degree, more explicitly, the experimental-specific artefact that may be not biologically relevant to translation. Therefore, our *MTtrans* framework is indeed important to allow multiple data types to be integrated to identify robust features that can work well across different conditions.

Although we have yielded robust regulatory motifs with maximally activated Seqlet (Figure 4), there is still rich information in the model that is not fully explained, such as the motif interaction. Recent studies analyse the motif interaction in an instance-based way. Ziga et al. identified the cooperative TF binding interaction by permuting the spacing of two candidate motifs [5]. Preserving the reading frame position of of the extracted feature is necessary for the model to correctly identify the in-frame or out-of-frame motifs [18]. For *MTtrans*, we choose to employ the GRU layers in the task-specific tower because it is effective at handling the spatial arrangement of the motif detected by the last convolutional layer. Visualisation of the GRU layer, however, usually requires an extra architectural plugin like attention [35]. Although the task-specific syntax explanation of the GRU layers is not the focus of this work, future work can reveal how the essential elements are coordinated to predict the translation rate.

Overall, *MTtrans* provides a solution to extract the robust translation regulatory elements in the 5’UTR from a collection of related yet noisy systems. We expect this multi-task framework can extend to learn the technique-invariant determinant for other sequence modelling questions to reveal the biologically meaningful signal from the sequence.

## Appendix

There is one additional file containing Supplementary Tables 1-2 and Supplementary Figures 1-8.

## Acknowledgements

We thank Dr. Sung Chul Kwon for his helpful suggestions about filtering the transcripts and Dr. Chen Qiao for help with model training. We also thank our colleague Xinyi Lin and Yiming Chao for their feedback on the data visualisation and testing the pipeline.

## Funding

This work was supported in part by AIRInnoHK administered by Innovation and Technology Commission and the National Natural Science Foundation of China Excellent Young Scientists Fund (32022089).

### Abbreviations

RP: Ribosome profiling;
MPRA: Massively Parallel reporter Assay;
FACS: Fluorescence-Activated Cell Sorting;
uAUG: upstream start codon;
uORF: upstream Open Reading Frame;
CNN: Convolutional Neural Network;
GRU: Gated Recurrent Unit;

## Availability of data and materials

The code to re-implement *MTtrans* can be access from https://github.com/holab-hku/MTtrans and the FACS library is also available are available from Gene Expression Omnibus (GEO) under the accession of GSE201766.

