## Supplementary Tables & Figures for "Translation rate prediction and regulatory motif discovery with multi-task learning"

### 1 Supplementary Tables

Supplementary Table. 1: Performance of different task combination strategies on each dataset. The rows are the consisting task, and the columns are different strategies. In each column of the table, a non-blank value showed the R-squared of MTtrans using the strategy on every test set and also noted which tasks were included in the combination. For example, strategy *M3R* included one MPRA dataset MPRA-H and all 3 RP datasets, so it only had values for these four rows.

| Combination | 3R | M3R | 2M3R | 3M3R | 3M2R | 3MR | 3M | single-task |
|---|---|---|---|---|---|---|---|---|
| MPRA-H | | 0.727 | 0.798 | 0.753 | 0.779 | 0.797 | 0.834 | 0.812 |
| MPRA-V | | | 0.807 | 0.82 | 0.83 | 0.845 | 0.871 | 0.864 |
| MPRA-U | | | | 0.883 | 0.892 | 0.907 | 0.942 | 0.941 |
| RP-muscle | 0.44 | 0.429 | 0.509 | 0.404 | | | | 0.34 |
| RP-pc3 | 0.456 | 0.443 | 0.499 | 0.471 | 0.468 | | | 0.39 |
| RP-293T | 0.364 | 0.378 | 0.379 | 0.396 | 0.38 | 0.317 | | 0.389 |

Supplementary Table. 2: The $p$-value of the statistical test for MTtrans has greater R-squared values. The one-sided unpaired T-test was performed with the alternative hypothesis of MTtrans has greater R-squared values than the two baselines.

| | mpra-u | mpra-h | mpra-v random | mprav-human |
|---|---|---|---|---|
| Mixing | 0.041559 | 0.000428 | 0.01496 | 0.781501 |
| FramePooling | 0.003711 | 0.008587 | 0.000194 | 0.003557 |

### 2 Supplementary Figures

### Additional Files

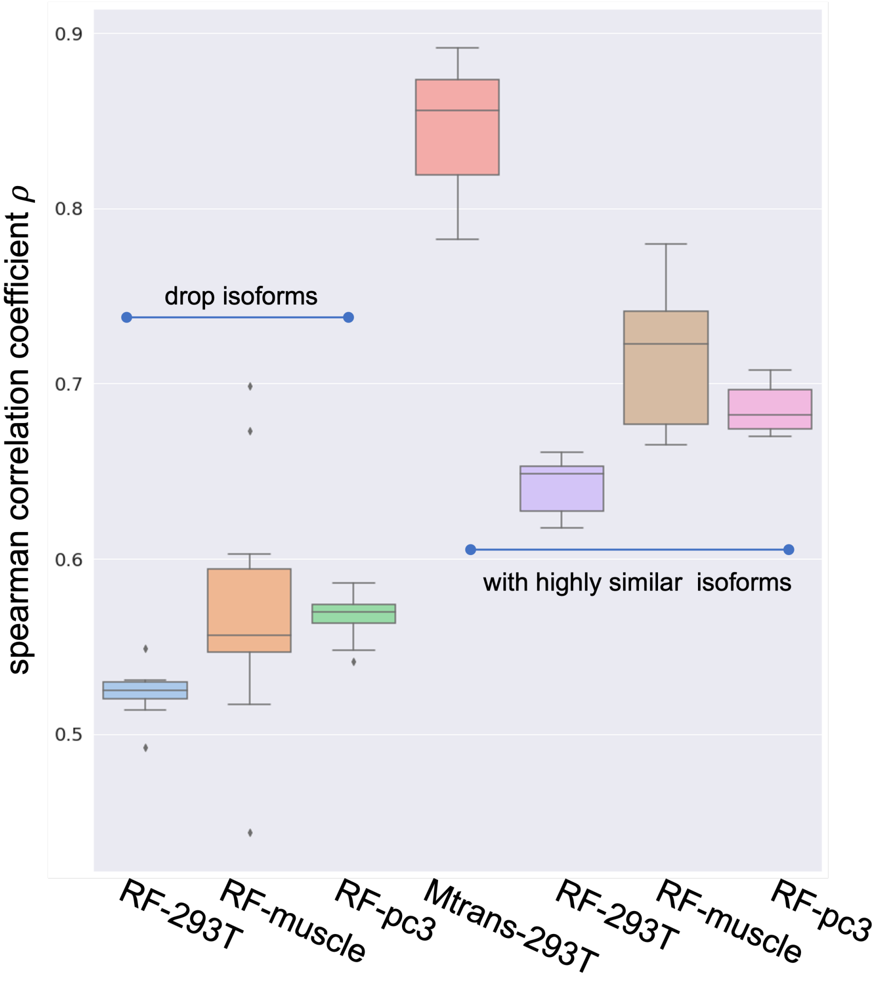


Supplementary Figure. 1: Data leakage from isoforms. The purpose of dropping out highly similar isoforms is to prevent the data leakage between training and testing set. The rightmost three models are the Random forest models trained from RP datasets without isoform selection and so does the *Mtrans* model in the middle. Deep learning based models benefits better than the RF models from the data leakage. After dropping isoform with identical last 100bp sequences, the prediction accuracy drops by around 15%, see the leftmost three.

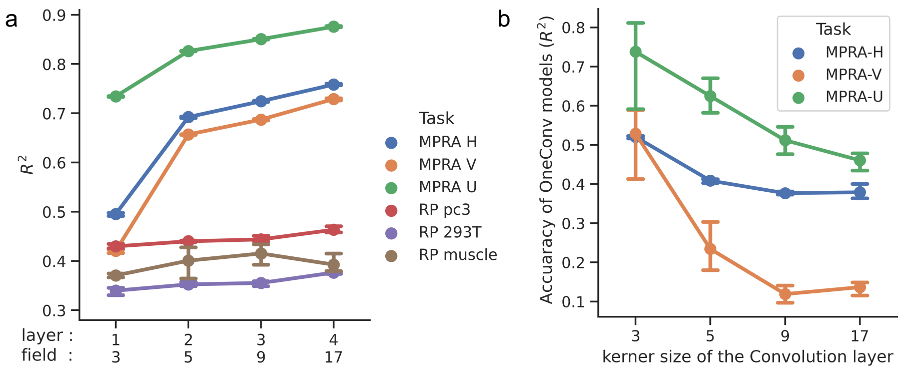


Supplementary Figure. 2: Interrogation of features from each layer.**a** The feature map encoded by each convolution layer of *MTtrans* was redirected as the input for random forest models, whose prediction accuracy indicates how informative each feature sets are. **b** The top 20 most important features for layer 2 and layer 3 features encoded by *3M3R*. The feature importance scores for the RF model trained for task RP-293T, RP-PC3 and MPA-H were sorted by their sum.

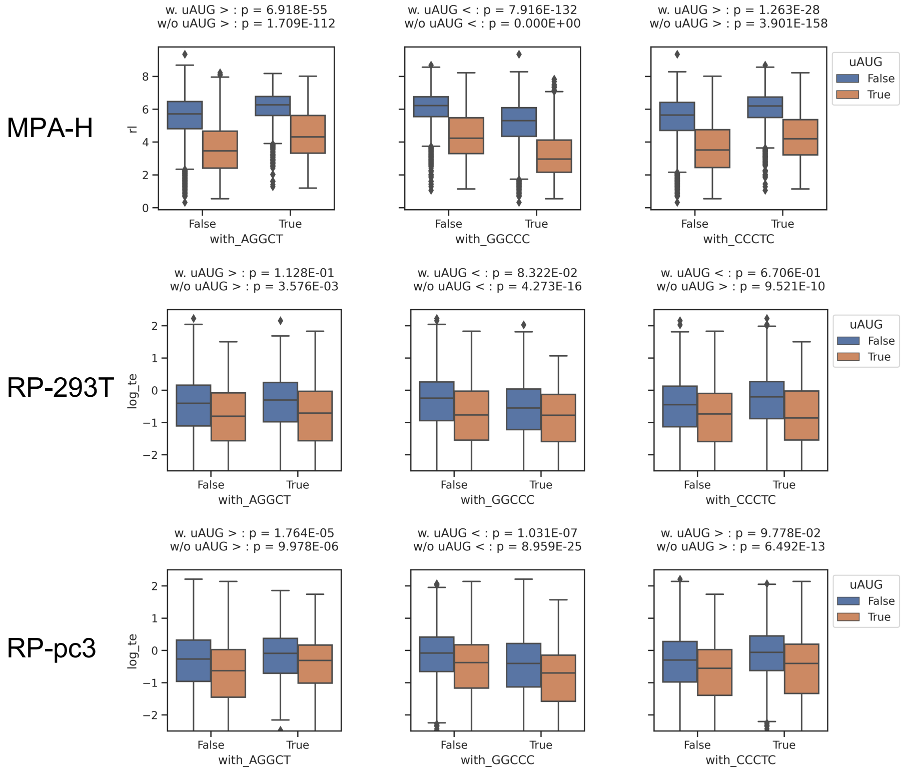


Supplementary Figure. 3: The effect of the top 3 motifs from laye2. The average translation rate, rl for task MPA-H and log(TE) for task RP-pc3 and RP-293T, is stratified by the co-occurrence of uAUG. *p*-value was calculated for the side with student t test, as both rl and log(TE) displayed the gaussian-like distribution.

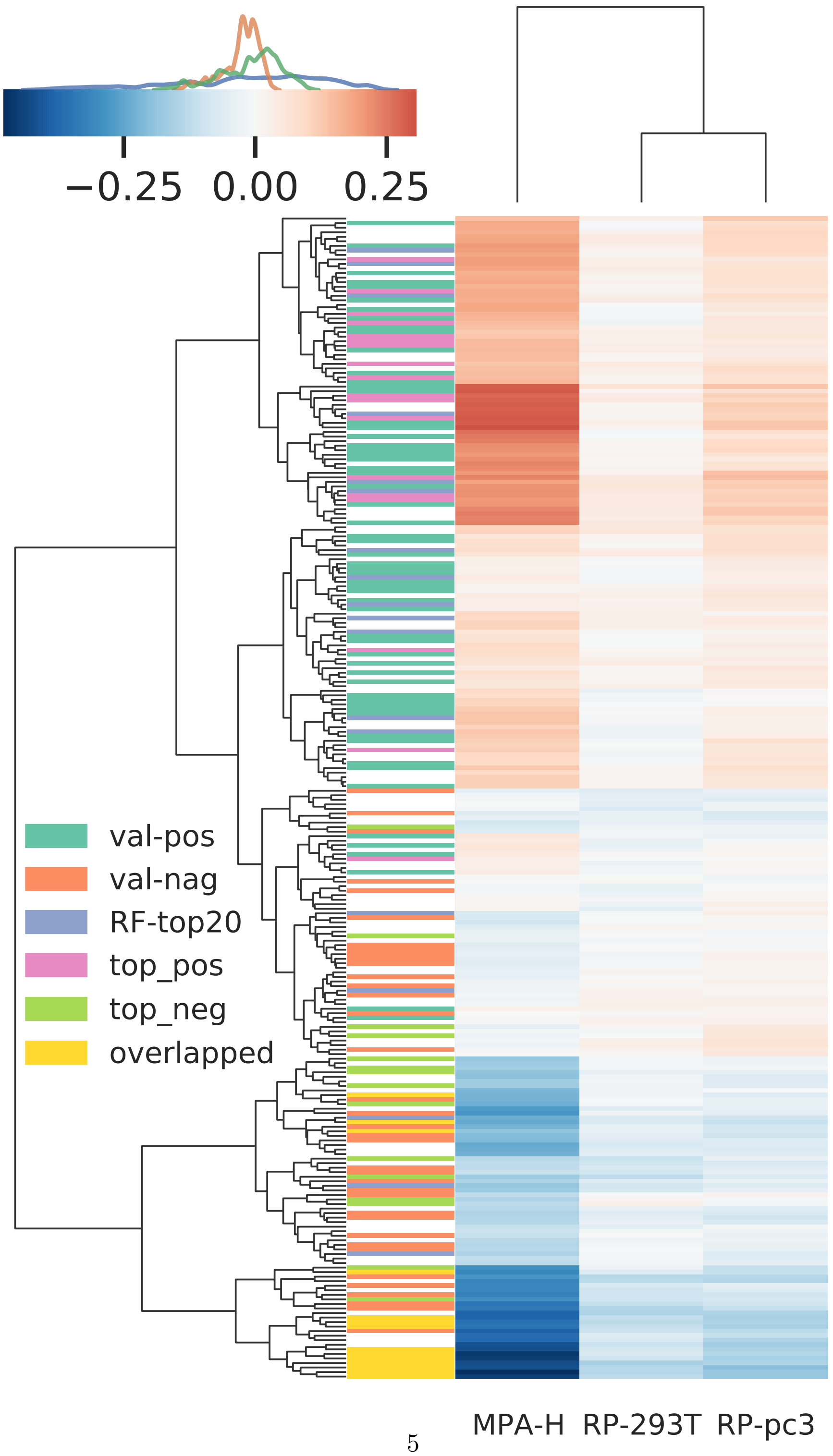




Supplementary Figure. 4: The heatmap of 256 discovered motifs. Row hierarchical clustering is applied to group motifs of similar regulatory effect (pearson correlation coefficient).

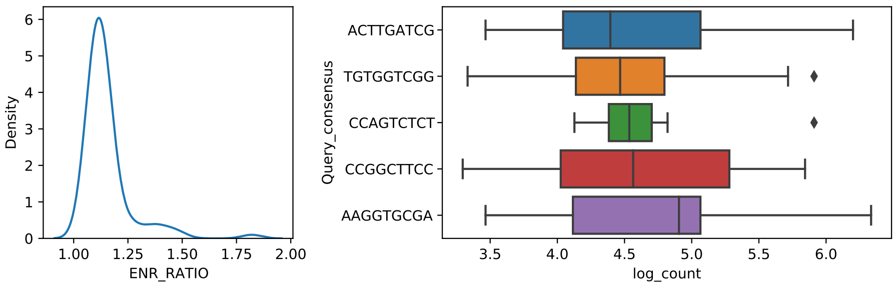


Supplementary Figure. 5: Motif enriched in the top 25% UTRs. 5 most enriched motifs are shown. The enrichment analysis was performed for all the positive motifs we found and searched in the 25% most abundant sequences, with the left 75% of the UTRs as the background

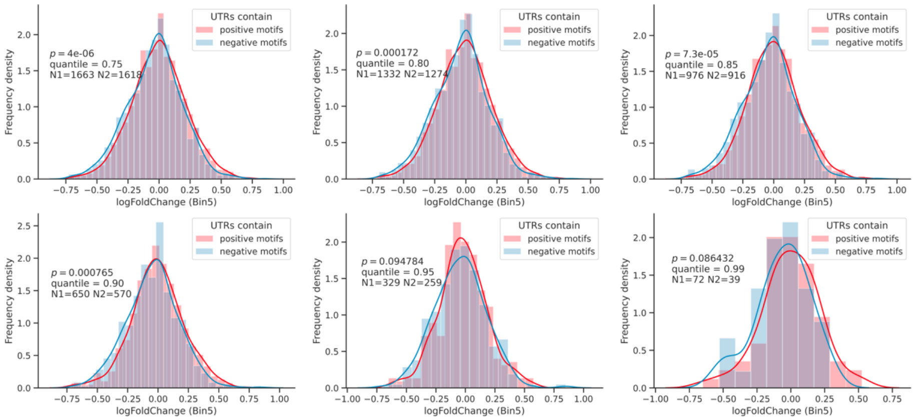


Supplementary Figure. 6: The distribution of translation rate for two shortlisted motif sets for quantile thresholds ranging from 0.75 to 0.99 in GSE176581. In each subplot, the positive group of UTRs (red) was selected from the library when the average PWM score of the 29 shortlisted positive motifs is higher than the defined quantile threshold. The negative group (blue) was formed in a similar way using the 25 shortlised negative motifs. N1 and N2 are the number of selected sequences in positive and negative group respectively. The $p$-values were obtained by performing one-sided unpaired T-test.

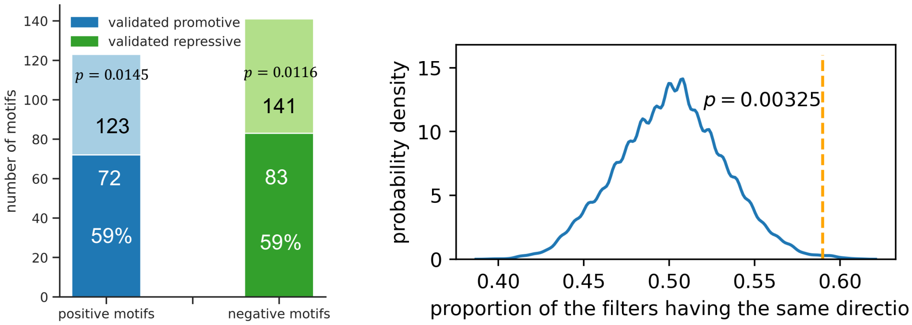


Supplementary Figure. 7: The proportion of motifs having consistent effect in our in-house FACS-seq library. **a**, out of 123 positive motifs (light blue), 72 of them (darker blue) could be validated to bring a higher translation rate than average by our independent FACS experiment (59%, p=0.0145). A oneside binomial test was conducted to examine how much this level is higher than random. Out of 141 negative motifs (light green), 83 of them (darker green) were also validated to be consistent (59%, p=0.0116). The $p$-value of testing the overall concordance was 0.0066 (153 out of 256). **b**, by scrambling the effect of motifs for 4,000 times, we generated the distribution of concordance level. The $p$-value is calculated using the number of observation having higher concordance proportion than 59%.

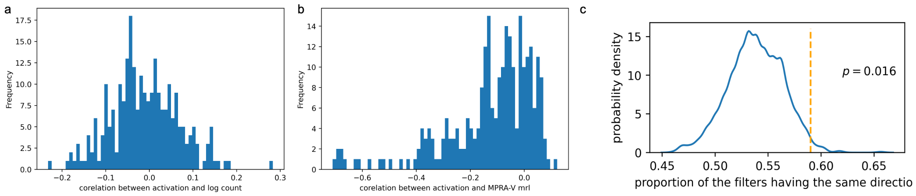


Supplementary Figure. 8: The Pearson correlation coefficient of convolutional filter activation and the translation rate. **a**, the distribution of Pearson correlation coefficient between activation value and log count in our in-house FACS-seq dataset. **b**, the distribution for MPRA-V dataset. **c**, The resampling test result for the concordance between coefficient learnt by logistic regression and correlation of MPRA-V dataset.
